# Zebrafish shoals share leadership during continuous decision-making on the move in a three-room Y-maze

**DOI:** 10.64898/2026.03.20.713221

**Authors:** Axel Séguret, Yohann Chemtob, Bertrand Collignon, Léo Cazenille, José Halloy

## Abstract

Collective decision-making in animal groups is often studied using short, trial-based mazes experimental setups that restrict observations to isolated choice events. However, how leadership and decision dynamics unfold over extended periods in symmetric environments remains poorly understood. Here we introduce a novel cyclic three-room Y-shaped environment that enables continuous, and autonomous sequences of collective decisions without experimental reset. We tracked the positions and identities of 20 groups of five AB-strand zebrafish (*Danio rerio*) during one-hour sessions in which animals freely transitioned between three identical rooms connected by visually isolated identical corridors. We show that this symmetric *Y-maze* enables the collection of large amounts of data to study decision-making with a few replicates, because habituation occurs after 45 minutes of exploration. After an initial exploration phase, groups reached a stable behavioural regime, generating thousands of decision events per replicate. Collective dynamics were consistent across spatial contexts, indicating that the symmetric architecture does not bias movement patterns, as opposed to traditional mazes. We show that zebrafish leadership is typically shared among shoal members, with leaders often acting as decision-makers. By transforming a classical maze into a self-renewing decision system, this approach enables the study of long-term collective dynamics and spontaneous leadership in controlled yet ecologically relevant conditions.

**Author summary:** présentation

## Introduction

Studies of animal collective behavior aim to understand and model inter-individual interactions and dynamics of social species at group levels. Shoals, schools, swarms, herds, flocks or stampedes are temporary and complex social organizations whose characteristics are influenced by the interactions between the individuals themselves and the environment [1–8]. Such hub-like (agora-like) collective structures facilitate conspecific encounters for a variety of tasks: information transfer, reproduction, nest site selection [9–12], migration, predator avoidance, food collection [13], etc. Depending on the species, these tasks rely on the collective decision-making of a few individuals (e.g. the personal leadership of mountain gorillas [14] or the distributed leadership of three-spined stickleback [15]), or the whole population (e.g. the consensus of honey bees [16], ants [17] or golden shiners [18]).

In ethology, many constrained environments have been developed to study decision-making, including corridor systems [5, 19, 20], Y-mazes [21], T-mazes [22], +-mazes [23, 24], radial mazes [25], or multi-door decision-making arenas, etc. These environments are simple to build, very versatile, and have provided valuable insights into pre-departure dynamics [26, 27], initiation [26, 28, 29], leadership [27, 28], follower organizations [28], or exploration [15, 30, 31]. These constrained experimental setups require the animals to travel alone or in a group through corridors or from sites to sites. For example, *Ward et al*. [21] have tested the preference of mosquitofish, *Gambusia holbrooki*, for corridors with or without a predator, or of damselfish *Dascyllus aruanus* for a piece of coral or a strip of coral sand and pebbles [28]. *Miller et al*. [24] have shown that mixed societies of golden shiners, *Notemigonus crysoleucas*, which have been trained through associative learning to choose one of two alternatives, will choose a third corridor that compromises the other two. During the trials, the animals are released in the setup, they move through the corridor to reach the decision area (generally the centre of the *Y, T* or *+*) where they choose a new corridor and a final site.

However, most of these setups share a common structural property: they are organized as discrete trials centered on a single decision event. Animals are typically released, make a binary or multi-arm choice, and are then removed for a few days [21, 24, 28] or repositioned in front the decision area after a few hours by experimenters [24, 32]. As summarized in Table 1, classical maze architectures often provide direct visual access to all alternatives from the decision point, lack stable residence zones within arms, and require human intervention between trials. This design constrains observations to short-term choice events and limits the study of endogenous sequences of collective decisions unfolding over extended time scales. Moreover, the constant human monitoring increases the amount of interactions between the studied animals and the humans, possibly biasing the experiments.

**Table 1.** Conceptual comparison of constrained decision-making environments. Classical maze paradigms are typically organized as discrete trials with a single decision event, often requiring human repositioning of animals between replicates. In contrast, the cyclic three-room Y-maze enables continuous autonomous sequences of symmetric collective decisions without experimental reset.

| Maze type | Typical decision structure | Visual access to alternatives | Residence zones | Human reset required | Suitable for long-term continuous dynamics | Leadership emergence detectable |
| --- | --- | --- | --- | --- | --- | --- |
| Corridor / two-area system | Single or binary decision per trial | Often visible | Yes (2 areas) | Yes | Limited | Moderate |
| Y-maze (classical) | Binary choice per trial | Both arms visible from center | No (arms are transit) | Yes | Low | Short-term only |
| T-maze | Binary choice per trial | Both arms visible | No | Yes | Low | Short-term only |
| Plus-maze / radial maze | Multiple simultaneous options | All arms visible | No | Yes | Low-Moderate | Short-term only |
| *n*-door arena | Multiple options | Usually visible | Variable | Yes | Moderate | Short-term only |
| Cyclic three-room Y-maze (this study) | Endless sequence of symmetric decisions | Corridors block visual access | Yes (3 identical rooms) | No | High (continuous 1h+ replicates) | Yes (spontaneous, repeated transitions) |

To address these limitations, we designed a cyclic and symmetric three-room Y-maze that transforms the classical Y architecture into a continuous and autonomous decision system (Fig. 1). Unlike the standard Y-maze, this unbiased *Y-maze* does not require any starting or ending zones to be defined. Each of the three identical arms consists of a corridor and a chamber (that can serve as a temporary residence zone), joining together at the centre of the *Y*, the decision area or triangle of decision (Fig 1). Crucially, corridors are visually isolated from one another, preventing individuals from perceiving alternative options before reaching the central decision area. Once a group reaches a room, no experimental reset is required: the arrival room immediately becomes the new departure site, allowing animals to engage in an endless sequence of symmetric decisions.

**Fig 1.**
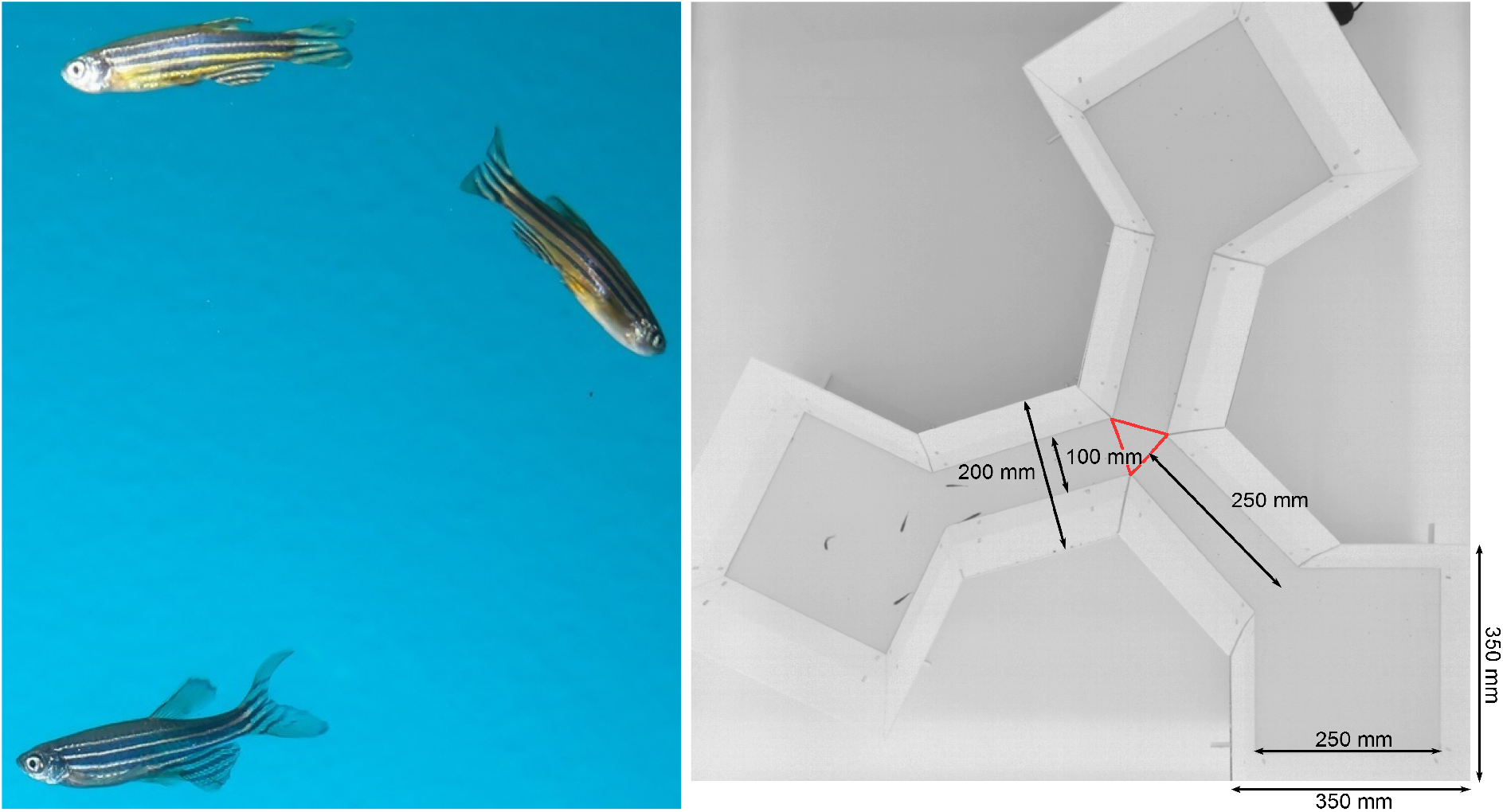
Experimental setup. **Left:** Three AB-strand zebrafish (*Danio rerio*). **Right**: the arena is composed by three square rooms (350 mm x 350 mm) connected by corridors (250 mm x 200 mm). We observed 20 groups of 5 zebrafish swimming during one hour to study their collective behaviours in a fragmented environment. The triangle of decision is drawn in red and superimposed on the picture.

In this study, we let the animals make successive choices between several identical alternatives (non-binary or symmetrical choices) in the 20 one-hour replicates. In the decision area, they can choose between the two rooms in front of them and the room from which they came. This set-up is a new variation of the previous one we built in [33, 34], *2 rooms connected by a corridor*, and always evokes patchy environments [35].

Here, we investigate whether a Y-maze connecting three rooms is suitable and unbiased for long-duration replicates in studies investigating the influence of leadership and collective decision-making among zebrafish. In particular, we aim to characterize the dynamics of the choices and the collective movements for a population size of 5 AB zebrafish in a constrained environment. Zebrafish are gregarious vertebrate model organisms used in behavioral [36, 37] and learning studies [38, 39]. They behave in groups in nature as much as in the laboratory [5, 40–42]. The size of the groups range from a few individuals to hundreds [43–45] depending on the region, the quality of the water (pH, temperature, turbidity, etc.) and the presence of predators, in a wide variability of habitats with varying structural complexities [45, 46].

## Results

### Habituation and biases of the setup

Collective decision metrics were computed across 20 replicates of groups of 5 AB zebrafish swimming in the Y-maze (Fig 1), without human interruption during one hour. We compute for four time intervals of 15 minutes the number of majority transitions between the two rooms (Fig 2). We count during the first 15 minutes of all the 20 replicates 1161 events of transitions (mean = 58.1, std = 13.5), for the second interval ([15; 30]) 1052 events (mean = 52.6, std = 15.5), for the third interval ([30; 45]) 976 events (mean = 48.8, std = 15.5) and for the last interval ([45; 60]) 850 events (mean = 42.5, std = 14.5). This corresponds to on average 202 majority decision events per trial in the twenty one-hour trials. The number of transitions decreases through time, and a Kruskal-Wallis test shows significant differences between the number of transitions for each interval (*p* − *value* < 0.01, *H* = 12.4 and *df* = 3). The results are significantly different: *p* − *value* < 0.01 between the intervals [0; 15] and [45; 60]. It means that the setup shows a low habituation rate within the experimental duration.

**Fig 2.**
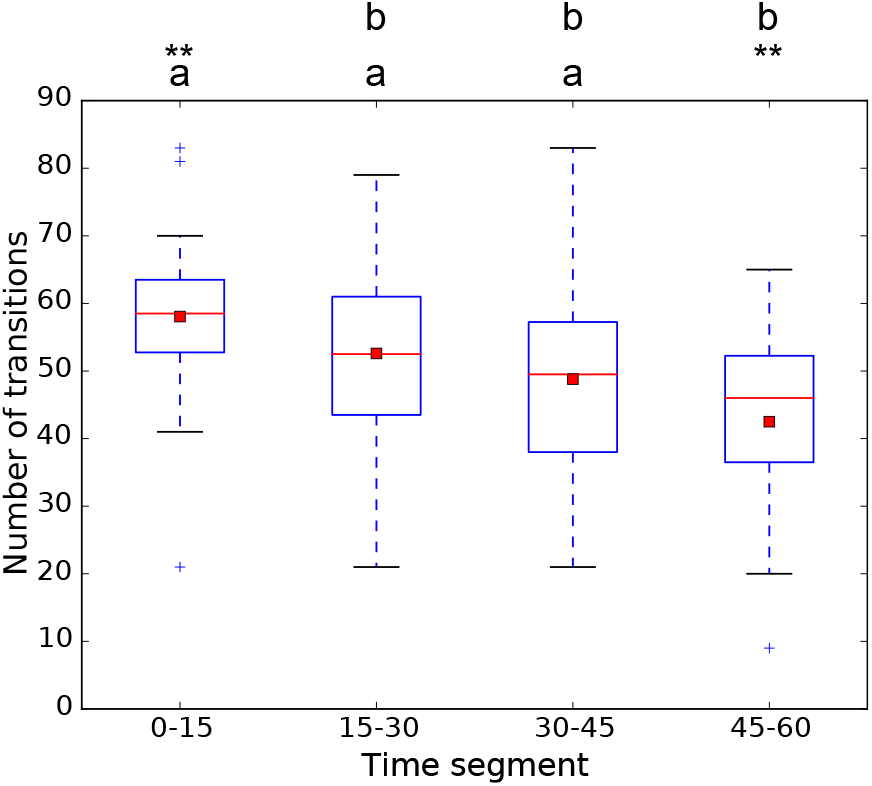
Number of majority transitions between two rooms at different steps of the experiment for 20 replicates with 5 zebrafish. The red line shows the median and the red square is the mean. * = *p* − *value* < 0.05, ** = *p* − *value* < 0.01, *** = *p* − *value* < 0.001, ns = non significant.

Then we focus on the transitions for groups of 4 or more zebrafish (Fig 3) to see if the setup shows any structural impact on the collective decision-making processes. We reveal that most of the time groups of zebrafish make decisions for the alternatives located in front of them (left or right): the mean of the probability for zebrafish to go to the left is 41.3%, to the right is 42.7% and to make a U-turn is 16.0%. The group transitions to the left and to the right rooms are significantly higher than U-turns: Kruskal-Wallis, *p* − *value* < 0.001, *H* = 34.1 and *df* = 2. A Tukey’s honest significant difference criterion, *p* − *value* < 0.001 for U-turns versus Left, and U-turns versus Right show significant differences.

**Fig 3.**
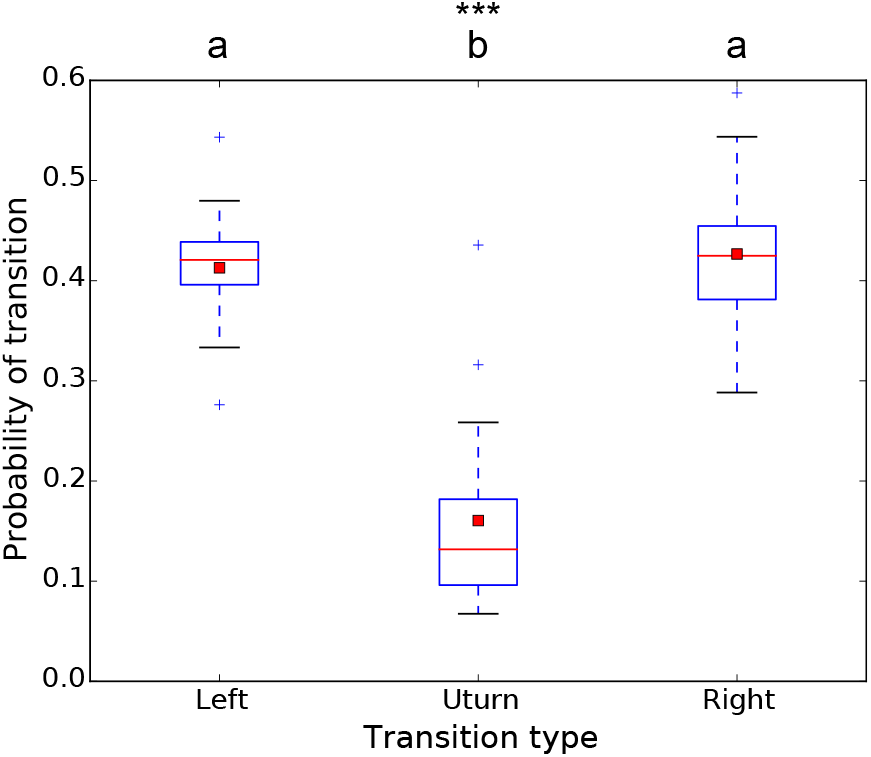
Transition probability for each fish to choose Left, Right or U-turn during a transition event of the majority of the population (⩾4 individuals). The red line shows the median and the red square is the mean. Zebrafish show the same probabilities to turn Left or Right and a lower probability to perform a U-turn. * = *p* − *value* < 0.05, ** = *p* − *value* < 0.01, *** = *p* − *value* < 0.001, ns = non significant.

### Collective departures and collective decision-making

In this section, we characterize initiation events, collective departure events (exits from any room) and collective decision events (exits from the triangle of decision, which is at the junction of the corridors). In addition, we focus on the identities of the fish in order to identify in each group temporary *leaders* and *decision-makers*. Thanks to idTracker software [47], we are able to identify and follow each fish. We consider a leader as the first zebrafish that leaves a room (initiation) and is followed by all the population (collective departure).

In the Fig 4, we show a strong correlation between the rank of exit of a room and the rank of exit of the triangle of decision. Also, this figure shows that, during each transition event, the leader and the decision maker generally are the same animal.

**Fig 4.**
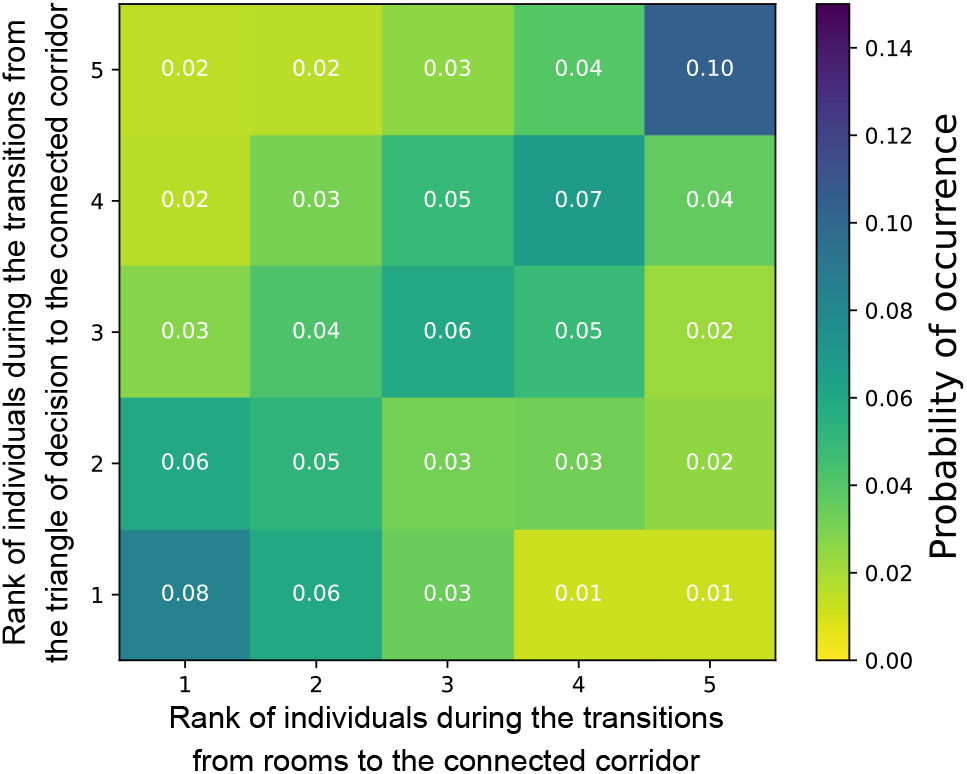
Change of ranks between exiting a room and exiting the triangle of decision. The first fish to exit the room is generally the first to exit the triangle of decision, i.e. the first to make a decision.

In the Fig 5, we reveal for the 20 replicates (A to T) the proportion of collective departures initiated by each fish and we rank the groups according to the score of the member that led with the highest proportion of departure. There is no replicate where the distributions of the initiations are equally shared by the zebrafish. In 25% of the replicates (P to T), the results show that one fish takes the position of the leader for at least 50% of the initiations.

**Fig 5.**
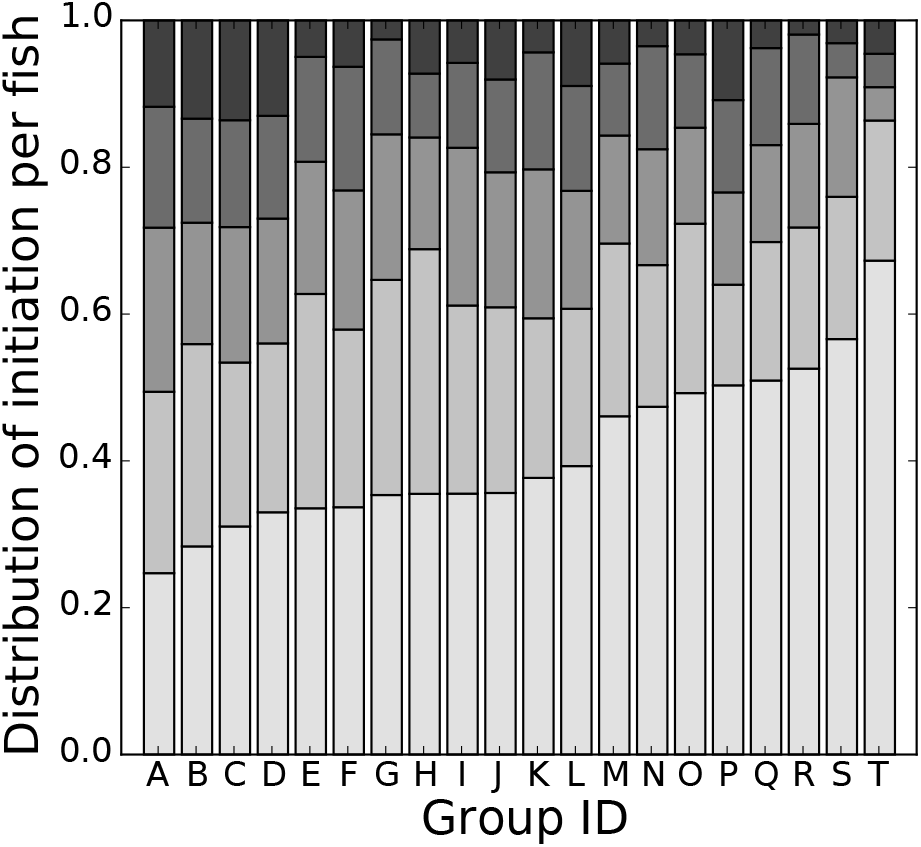
Proportion of collective departures. led by each group member. The fish are ranked in each group according to the proportion of initiation that they made and the groups are ranked according to the highest proportion of departure shown by one of a group member. The leadership is unequally shared between individuals. In 5 groups, one individual leads the group for at least 50% of the initiations.

In the Fig 6, for all replicates we look at the changes of leadership over time, according to 4 time intervals, between consecutive events of decision in the triangle of decision. We define the change of leadership as a change of initiator between two simultaneous initiations. There are 957 values for the interval 0 to 15 minutes, 827 values for the interval 15 to 30 minutes, 721 values for the interval 30 to 45 minutes, and 641 values for the interval 45 to 60 minutes. The mean of the proportions of changes in leadership oscillate around 70% regardless of the time interval. A Kruskal-Wallis test shows that regardless of the time interval, there are no significant differences between the distributions in the changes in the leadership (*p* − *value* > 0.05, *H* = 14.1 and *df* = 3). The same kind of analysis has been done between consecutive events of decision at the exit of a room (Fig S9). The results are similar.

**Fig 6.**
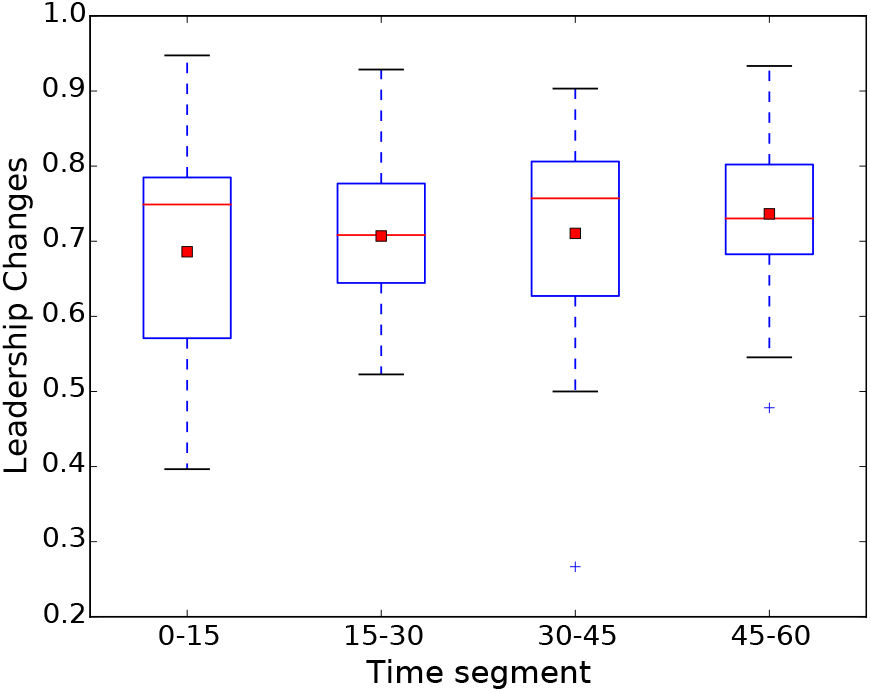
The leadership changes at each transition with a high probability and during the whole duration of each replicate. We compared the changes of leadership between consecutive events of decision in the triangle of decision. The red line shows the median and the red square is the mean. No significant differences between the distributions in the changes in the leadership were observed (*p* − *value* > 0.05, *H* = 14.1 and *df* = 3).

## Discussion

We study, over one-hour-long replicates, the dynamics of the leadership and the decision making of 20 different groups of 5 AB naive zebrafish in an unbiased Y-maze connecting three identical resting areas through three identical corridors. In this study, we show that our setup is unbiased: Fig 3 does not show significant differences in the preference of zebrafish groups to choose one of the two front arms (left or right) when coming from the third arm.

We already know that the personal motivation of one animal can initiate collective departures. They have been studied in many species: *Equus burchellii* [48], *Dicentrarchus labrax* [49], or *Gasterosteus aculeatus* [50]. Depending on the species, before collective departures, the location of the initiator (called leader when the collective departure is successful) within the group is of great importance. While for monkeys (*Cebus capucinus* or *Macaca mulatta*), the individuals in the center of the group generally initiate collective departures [51, 52], for fish (*Notemigonus crysoleucas, Rutilus rutilus*, or *Gasterosteus aculeatus*), this role is attributed to the animals on the periphery of the group [29, 53–55]. In addition, bolder individuals have been investigated to be predisposed to leadership in a variety of species of fish and birds (*Notemigonus crysoleucas, Branta leucopsis* and *Lepomis macrochirus*) [56–58]. In zebrafish, the authors also show that personal motivation is related to the intrinsic propensity to explore a new environment [12, 59]. This means that potential leaders choose their positions in the shoals in order to maximize the likelihood of a successful collective departure.

We build our setup to investigate leadership and decision-making. In our study, among the 20 independent groups of 5 AB zebrafish studied, we reveal a wide spectrum of leadership types (Fig 5). In 15 groups (A to O), the leadership is totally to partially shared between individuals, when for 5 groups the leadership is not shared, as at least 50% of the initiations are performed by the same fish (P to T). Moreover, among the 100 zebrafish studied, 62 zebrafish led initiations less than 20% of the time. By computing the data about leadership, we reveal that a change in leadership is a common practice in zebrafish. The turnover of leadership remains high (∼ 70%) and does not significantly vary over time. We hypothesize that this high and stable turnover reflects the absence of a rigid dominance hierarchy in AB zebrafish, consistent with the egalitarian and short-lived leadership reported in freely swimming shoals [34, 60]. This rotation may reflect fluctuating individual motivational states (e.g., hunger, exploration drive) that transiently predispose different individuals to initiate movement [61], rather than a fixed leadership role. Such distributed leadership may serve an adaptive function by sharing the energetic costs and predation risks of leading among group members [62]. We also wonder if groups P to T were composed of four shy individuals and one bold individual, or if it exists a spectrum of individual singularities: for example the boldness of those leaders could be stronger than the boldness of the other fish. Revealing individual singularities in shoals requires long-term observations (days to weeks). We are currently developing a new protocol for measuring these singularities, which will enable us to collect sufficient data on each individual. This will be the topic of another publication.

After the animals engage in a corridor, they reach the junction of the three arms (triangle of decision), where decision-making should occur. Here, they have the choice between the right arms and the left arms, or to move to the previous known location (U-turns). Our findings show that collective U-turns are peculiar events. They occur at the expense of losing speed and a reorganization of the group [63]. U-turns in zebrafish have been studied according to various structural complexities: from simple landmarks in an open environment [64] to networks of two [33] or three rooms (this study) as different parts of constrained environments. Our new constrained environment tends to replicate the natural environment of zebrafish (from river channels, irrigation canals, to beels) [45, 46]. AB zebrafish groups seem to favor collective movements to new visible locations over collective U-turns, regardless of the structural complexity (landmarks, networks of corridors and rooms). This consistent behavioral preference strengthens collective exploration, thereby increasing the group’s opportunity to access potential food sources with less loss of speed and few group reorganizations. These hypotheses are strengthened by the positive correlation between the role of leader and the role of decision maker shown in Fig 4.

Our findings on leadership dynamics in zebrafish contrast sharply with those reported for mosquitofish (*Gambusia holbrooki*), providing insight into how social structure shapes collective decision-making across species. *Burns et al*. [65] tested 18 populations of 5 Mosquitofish in a Y-maze over 5 replicates per population (one replicate per day). Once the fish made their decision they were captured and put back in their breeding tank. Unlike *Gambusia holbrooki* in a Y-maze where the “leadership switched between group members more often in an unfamiliar environment than when group members had experience of that same environment”, the leadership of zebrafish does not evolve over time (Fig 6). In the literature, their experiments were the closest to ours. Both of our teams tested the effect of unfamiliar and familiar environments on leadership. In our experiments, we consider the unfamiliar environment as the first minutes of the replicates, after a period of habituation [64]. We suggest that this contradiction between the leaderships in groups of zebrafish and mosquitofish may be due to their propensities to grasp their territory and to deal with the group. It seems that zebrafish and mosquitofish have developed opposite adaptive collective movements. Mosquitofish are known to form highly hierarchical societies [66, 67] when our observations suggest that zebrafish share the leadership equally to partially equally (Figures 5 and 6). This hypothesis is strengthened by Figures S8 and S9: there is no clear leader over a long period of time, and each fish can initiate a collective departure. We confirm the hypothesis we developed in [34].

In addition, we express reservations about the way experiments in mazes, generally based on repetitive tasks and interrupted by humans, are carried out and the meaning of these results (Table 2). We think that the animals should be naive for such maze experiments. At each replicate in the maze, the animals experience stress, endure human interactions, and may partially learn the experimental area, since each time they make a choice or after a time of exploration, they are returned to their breeding tank which is also less stressful. If the animals are not naive, it is therefore possible that (a) the group reproduces the same behavior as in the previous replicate and (b) the individuals who led the group during the previous replicate may reproduce this role and reinforce it. If the quantification of group features, such as travel time between rooms or collective transitions, change during the one-hour replicate, this suggests that the animals may be becoming familiar with the setup. Figures 2 and S5 show that the numbers of transitions, and of majority events in the corridors and in the rooms decrease with time when the travel time through the corridors increases slowly over the four time intervals. We show a significant difference between the first time interval [0-15] and the last [45-60]. As our results seem to show that the setup is unbiased, we can conclude that the animals become familiar with the setup after 45 minutes. By reducing human intervention and allowing the fish to swim freely for an extended period, it is possible to minimize stress, reduce bias in collective movement and decision-making, while demonstrating limited environmental learning.

**Table 2.** Use of mazes for recent decision-making studies with fish. min for minutes, sec for seconds, * replicates with 1 fish, ** replicates with 2 and 4 fish, *** replicates with 8 and 16 fish. Disambiguation: *Duration of each replicate* means the duration between the beginning of the replicate (release of the animals) and the end of the replicate (when the animals reached an arm or when the experimenter decided to stop the record).

| Authors | Type of maze | Number of replicates | Number of events of decision-making | Duration of each replicate | Number of replicates per animal | Length of the corridors |
| --- | --- | --- | --- | --- | --- | --- |
| Ward et al. [68] | Y-maze | 320 | 320 | few min | unknown or 20 times | two corridors of 75 cm |
| Sumpter et al. [69] | Y-maze | 2200 | 2200 | few min | unknown or 20 times | two corridors of 75 cm |
| Couzin et al. [18] | Two attractive areas | 108 | 108 | unknown | 3 times | 35 cm x 35 cm for each area in a 210 cm x 120 cm tank |
| Ward et al. [21] | Y-maze | 107 *<br>12 **<br>13 *** | 107<br>12<br>13 | few sec<br>few sec<br>few sec | unknown<br>unknown<br>unknown | two corridors of 50 cm and one of 80 cm |
| Burns et al. [65] | Y-maze | 90 | 90 | unknown | 5 times | two corridors of 50 cm and one of 80 cm |
| Ward et al. [70] | Y-maze | 240 | 240 | few min | only once | two corridors of 75 cm |
| Benhaim et al. [71] | T-maze like | 90 | 90 | 10 min | 3 times | three corridors of 20 cm connected to a 80 cm corridor |
| Miller et al. [24] | Y-maze like | 640 | 640 | 5 min | 40 times | six corridors of 28 cm |
| Woodley et al. [72] | Y-maze | 16 | unknown | 5 min | not known | four corridors of 46 cm |
| Ioannou et al. [73] | Y-maze | 102 | 4730 | 5 min | only once | three corridors of 36.5 cm |
| Delcourt et al. [25] | Radial-maze | unknown<br>9 | unknown<br>unknown | 60 min<br>720 min | unknown<br>unknown | six corridors of 42 cm |
| Camberlain et al. [74] | V-maze | 69 | 690 | 20 min | 10 times | two arms of 48 cm |
| Searle [75] | Y-maze | 38 | 38 | 10 min | only once | two corridors of 25 cm and one corridor of 27 cm |
| Li et al. [76] | Radial-maze | 64 | unknown | 20 min | only once | six corridors of 42 cm |
| **This study** | Y-maze | 20 | 4039 | 60 min | only once | three corridors of 25 cm connected to a room of 25 cm x 25 cm at each end |

The objective of the unbiased *Y-maze* is to achieve a balance between minimizing animal stress, reducing biases related to the setup and animal-human interactions while maintaining a low habituation rate. This ensures that the animals’ behavior is less influenced. Table 2, which summarizes recent studies using mazes, shows that it can take a while to perform all the replicates required to obtain sufficient data. We generally observe that the animals are studied several times for the same experiments, which seems to bring more biases. In the unbiased *Y-maze*, we allow the fish to swim freely over a long period of time, enabling us to collect a large amount of behavioral data while minimizing stress. For example, after one hour, each group took on average 202 decisions. In a few days, with minimal human interactions, it is possible to test 20 groups, showing more than 4000 collective decision events.

By converting a classical binary maze into a cyclic and self-renewing decision system, we provide a methodological framework that enables the study of long-term collective dynamics under controlled yet minimally intrusive conditions. This approach bridges the gap between short, trial-based paradigms and fully naturalistic observations. Beyond zebrafish, such cyclic architectures may serve as powerful tools to investigate distributed decision-making, leadership plasticity, and collective exploration strategies across social species.

## Methods

### Ethic statement

Fish experiments were performed in accordance with the recommendations and guidelines of the Buffon Ethical Committe (registered to the French National Ethical Committee for Animal Experiments #40) after submission to the state ethical board for animal experiments.

### Animals and housing

We bred 100 AB strain laboratory wild-type zebrafish (*Danio rerio*) up to the adult stage and raised them under the same conditions in tanks of 3.5L by groups of 20 fish in a zebrafish aquatic housing system (ZebTEC rack from Tecniplast) that controls water quality and renew 10% of water in the system every hour. Zebrafish descended from AB zebrafish from different research institutes in Paris (Institut Curie and Institut du Cerveau et de la Moelle Épinière). AB zebrafish show zebra skin patterns and have short tail and fins. They measure about 3.5 cm long. All zebrafish observed in this study were one year old at the time of the experiments. We kept fish under laboratory conditions: 27 ^*◦*^C, 500*µ*S salinity with a 10:14 day:night light cycle, pH is maintained at 7.5 and nitrites (NO^2−^) are below 0.3 mg/L. Zebrafish are fed two times a day (Special Diets Services SDS-400 Scientic Fish Food).

### Experimental set-up

The experimental tank (Fig 1) consists in a 1.2 m x 1.2 m tank confined in a 2 m x 2 m x 2.35 m experimental area surrounded by white sheets, in order to isolate the experiments and homogenise luminosity. The three squared rooms (350 mm x 350 mm) connected by corridors (250 mm x 200 mm) starting at one corner of each room (Fig. 1) have been designed on a Computer-Aided Design (CAD) software and cut out from Poly(methyl methacrylate) (PMMA) plates of 0.003m thickness. Each wall are titled, (20° from the vertical) to the outside with a vertical height of 0.14 m, to avoid the presence of blind zones for the camera placed at the vertical of the tank. The water depth is at 6 cm in order to keep the fish in a nearly two-dimensional space to facilitate their tracking, the pH is maintained at 7.5 and Nitrites (NO^2−^) are below 0.3 mg/L. A high resolution camera (2048 px x 2048 px, Basler Scout acA2040-25gm) is placed above the water surface (160 cm), at the vertical, and records the experiment at 15 fps (frame per second). The luminosity is ensured by 4 LED lamps of 33W (LED LP-500U, colour temperature: 5500 K - 6000 K) placed on each corner of the tank, above the aquarium and directed towards the walls to provide indirect lightning.

### Experimental procedure

We recorded the behavior of zebrafish swimming in the setup, during one hour, and did 20 replicates of groups of 5 naive AB zebrafish. Before each replicate, the starting room, from which the fish are released, is chosen randomly. Then, the fish were placed with a hand net in a cylindrical arena (20 cm diameter) made of plexiglas in the center of the selected room. Following a 15 minutes acclimatization period, the camera started recording, the fish were released and free to swim in the set-up. After one hour, the record stops and the fish are caught by a hand net and replaced in the rearing facilities. The fish were randomly selected regardless of their sex and each fish was never tested twice to prevent any form of learning.

### Tracking & data analysis

Today, the studies on animal collective behaviours use methodologies based on massive data gathering, for example for flies (*Drosophila melanogaster*) [77, 78], birds (*Sturnus vulgaris*) [79–81], fish (*Notemigonus crysoleucas*) [82]. Our experiments were tracked by post-processing (“off-line”) with the idTracker software to identify each fish and their positions [47]. This multitracking software extract specific characteristics of each individual and uses them to identify the animal throughout the video. This method avoids error propagation and is able to successfully solve crossing, superposition and occlusion problems. Each replicate consists of 54000 frames, corresponding to 270000 tracked positions (5 individuals over 54000 time steps): *P* (*x, y, t*) with coordinates *x* and *y*, and *t* the time.

We build the trajectories of each fish and compute individuals and collective measures (position in the arena, speed, acceleration). The instantaneous speed *v*_*t*_ was calculated on three positions and thus computed as the distance between *P* (*x, y, t* − 1) and *P* (*x, y, t* + 1) divided by 2 time steps while the instantaneous acceleration was computed as Δ*s/*Δ*t*.

## Authors’ contributions

AS carried out the lab work, the data analysis and the design of the set-up and drafted the manuscript; YC carried out the lab work, the design of the set-up, the data analysis and drafted the manuscript; BC carried out the statistical analyses and participated in the data analyses and in the writing of the manuscript; LC developed the online tracker, stabilized the video acquisition and improved idTracker and helped write the manuscript; JH conceived the study, designed the study, coordinated the study and helped draft the manuscript. All authors gave final approval for publication.

## Acknowledgements

This work was supported by European Union Information and Communication Technologies project ASSISIbf, FP7-ICT-FET-601074. The funders had no role in study design, data collection and analysis, decision to publish, or preparation of the manuscript.

