## Supplementary Information for "Zebrafish shoals share leadership during continuous decision-making on the move in a three-room Y-maze"

**1** all as one, Paris, France
**2** Laboratory of Social Evolution and Behavior, The Rockefeller University, New York, NY, USA
**3** Solvay Brussels School of Economics & Management, Université libre de Bruxelles (ULB), Brussels, Belgium
**4** Sorbonne Université, CNRS, ISIR, F-75005 Paris, France

◑These authors contributed equally to this work.
¤Current Address: Dept/Program/Center, Institution Name, City, State, Country
* ** ***

### 1 Supporting information

Supplementary figures of "Zebrafish (*Danio rerio*) collective behaviours in an unbiased Y-maze connecting three areas".

#### 1.1 Habituation

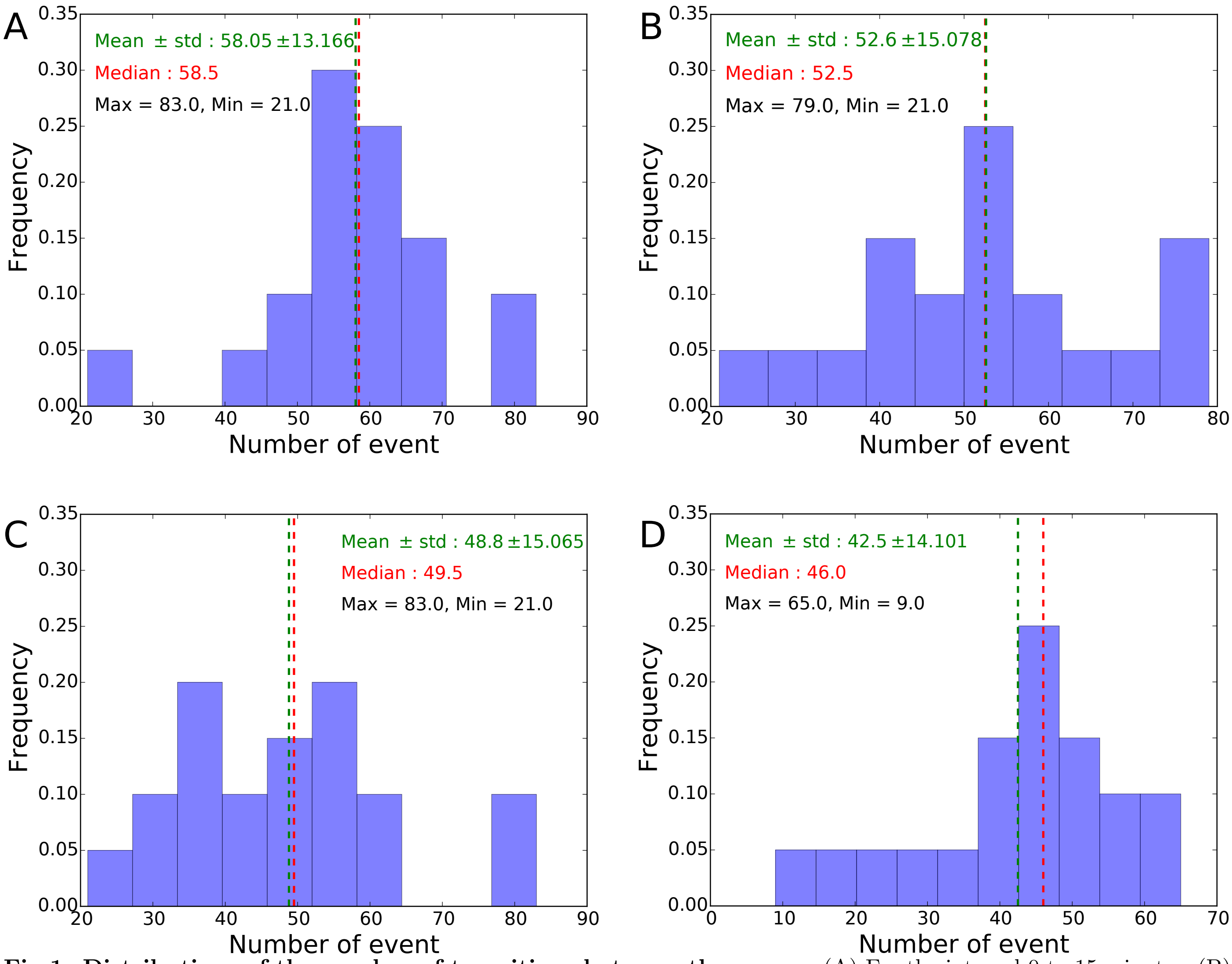


**Fig 1. Distributions of the number of transitions between the rooms**. (A) For the interval 0 to 15 minutes, (B) for the interval 15 to 30 minutes, (C) for the interval 30 to 45 minutes, (D) for the interval 45 to 60 minutes.

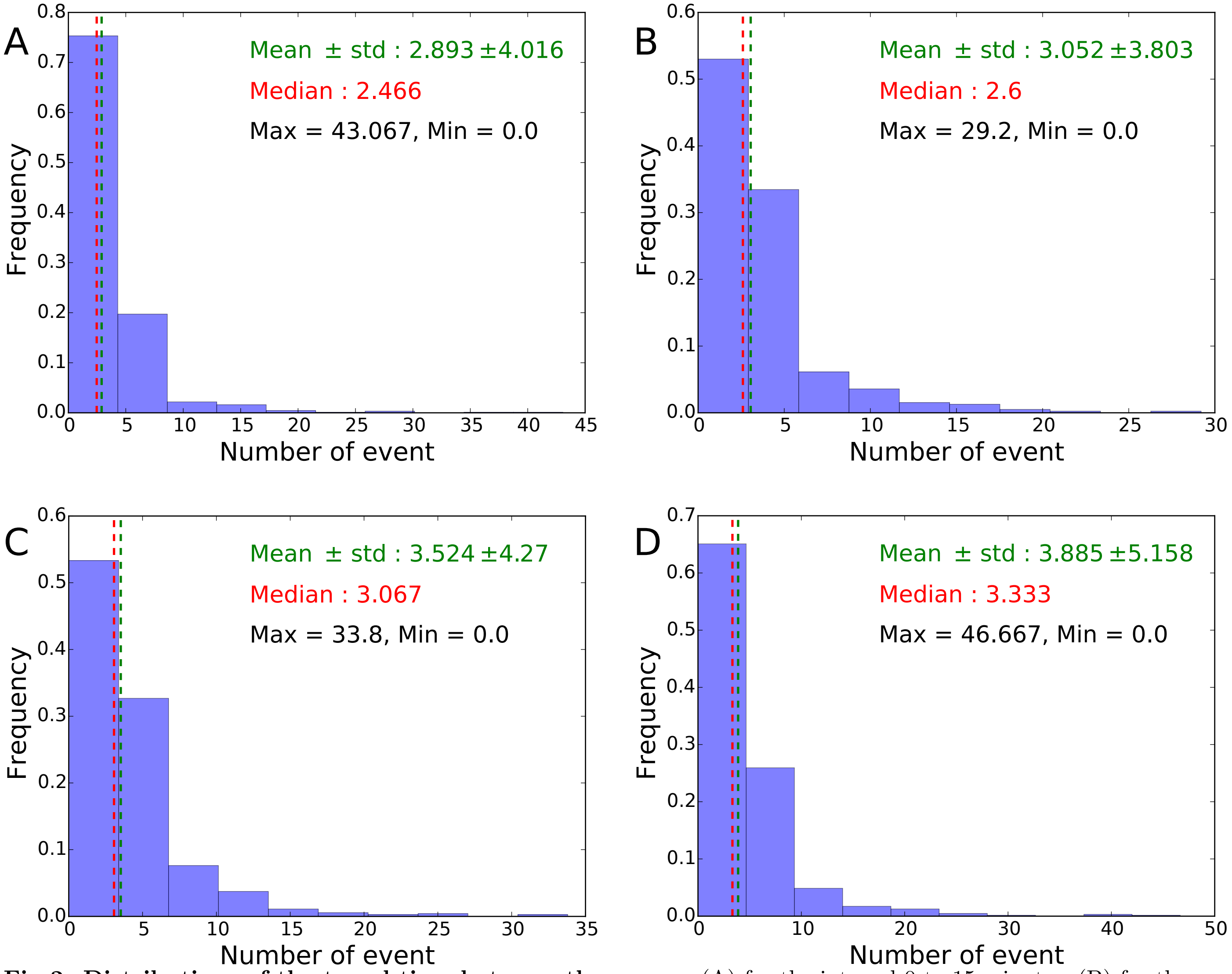


**Fig 2. Distributions of the travel time between the rooms**. (A) for the interval 0 to 15 minutes, (B) for the interval 15 to 30 minutes, (C) for the interval 30 to 45 minutes, (D) for the interval 45 to 60 minutes.

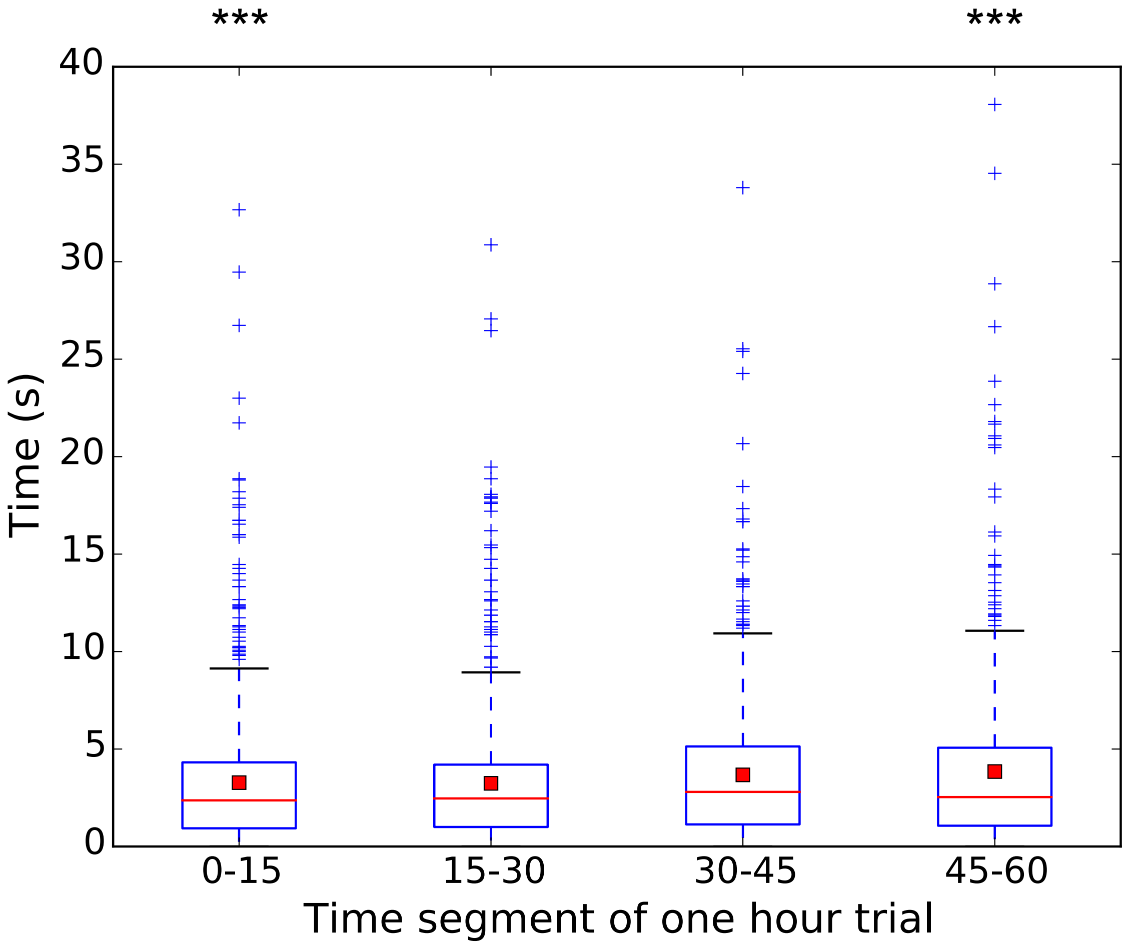


**Fig 3. Travel time between two rooms** at different steps of the experiment for 20 replicates with 5 zebrafish. The red line shows the median and the red square is the mean. A Kruskal-Wallis test shows no significant evolution during one hour of experiment ($p < 0.01$). $* = p - value < 0.05$, $** = p - value < 0.01$ and $*** = p - value < 0.001$.

We compute for four time intervals of 15 minutes the travel time beetween the rooms (Fig 3). The first interval is based on 867 values, the second on 783 values, the third on 722 values and the last one on 636 values. A Kruskal-Wallis test shows that there are significant differences between the time travels between the intervals ($p - value < 0.001$, $H = 22.5$ and $df = 3$). A Tukey's honest significant difference criterion shows that $p - value < 0.001$ between the intervals [0; 15] and [45; 60].

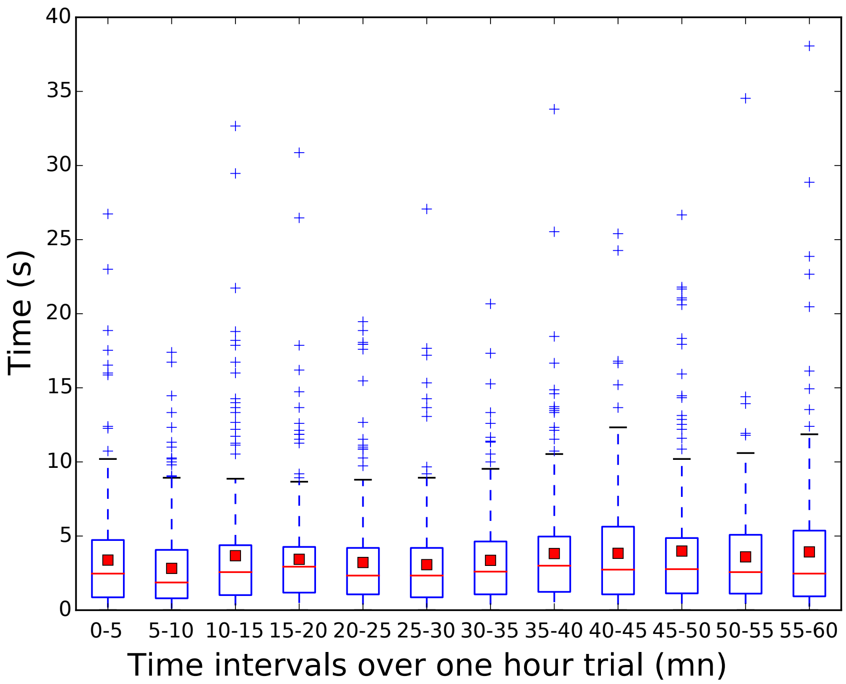


**Fig 4. Travel time between two rooms** at different steps of the experiment for 20 replicates with 5 zebrafish. The red line shows the median and the red square is the mean. A Kruskal-Wallis test shows no significant evolution during one hour of experiment ($p > 0.05$).

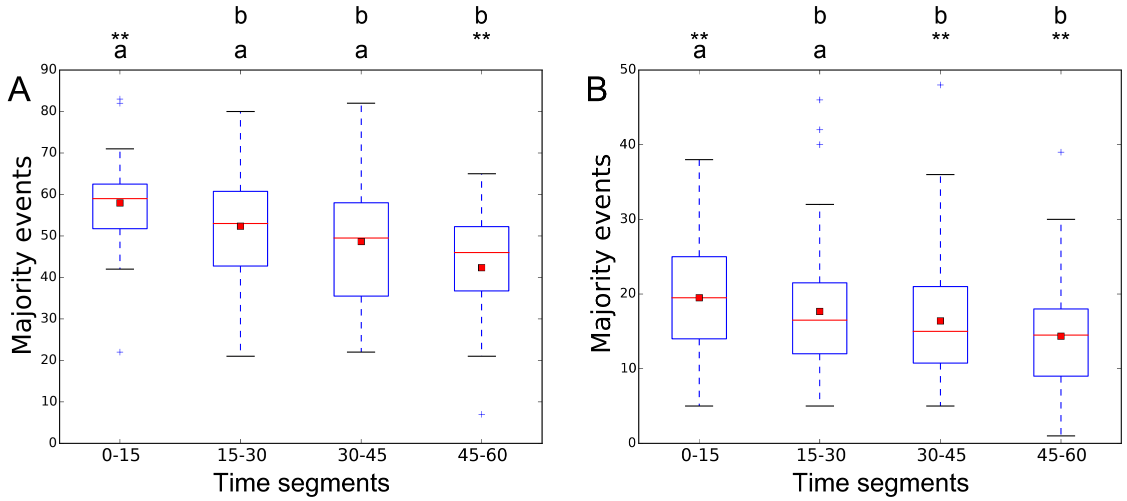


**Fig 5. Number of majority events** at different steps of the experiment for 20 replicates with 5 zebrafish. (A) in the corridors, (B) in the rooms. The red line shows the median and the red square is the mean. (A) The number of majority events in the corridor are significantly higherfor the time interval [0, 15] than [45, 60] (Kruskal-Wallis, $p - value < 0.01$, Tukey's honest significant difference criterion, $p - value < 0.01$ for[0, 15] versus [45, 60]). (B) The number of majority events in the corridor are significantly higherfor the time interval [0, 15] than [45, 60] (Kruskal-Wallis, $p - value < 0.01$, Tukey's honest significant difference criterion, $p - value < 0.01$ for[0, 15] versus [30, 45] and for[0, 15] versus [45, 60]). * $= p - value < 0.05$, ** $= p - value < 0.01$, *** $= p - value < 0.001$, ns = non significant.

### 1.2 Collective decisions

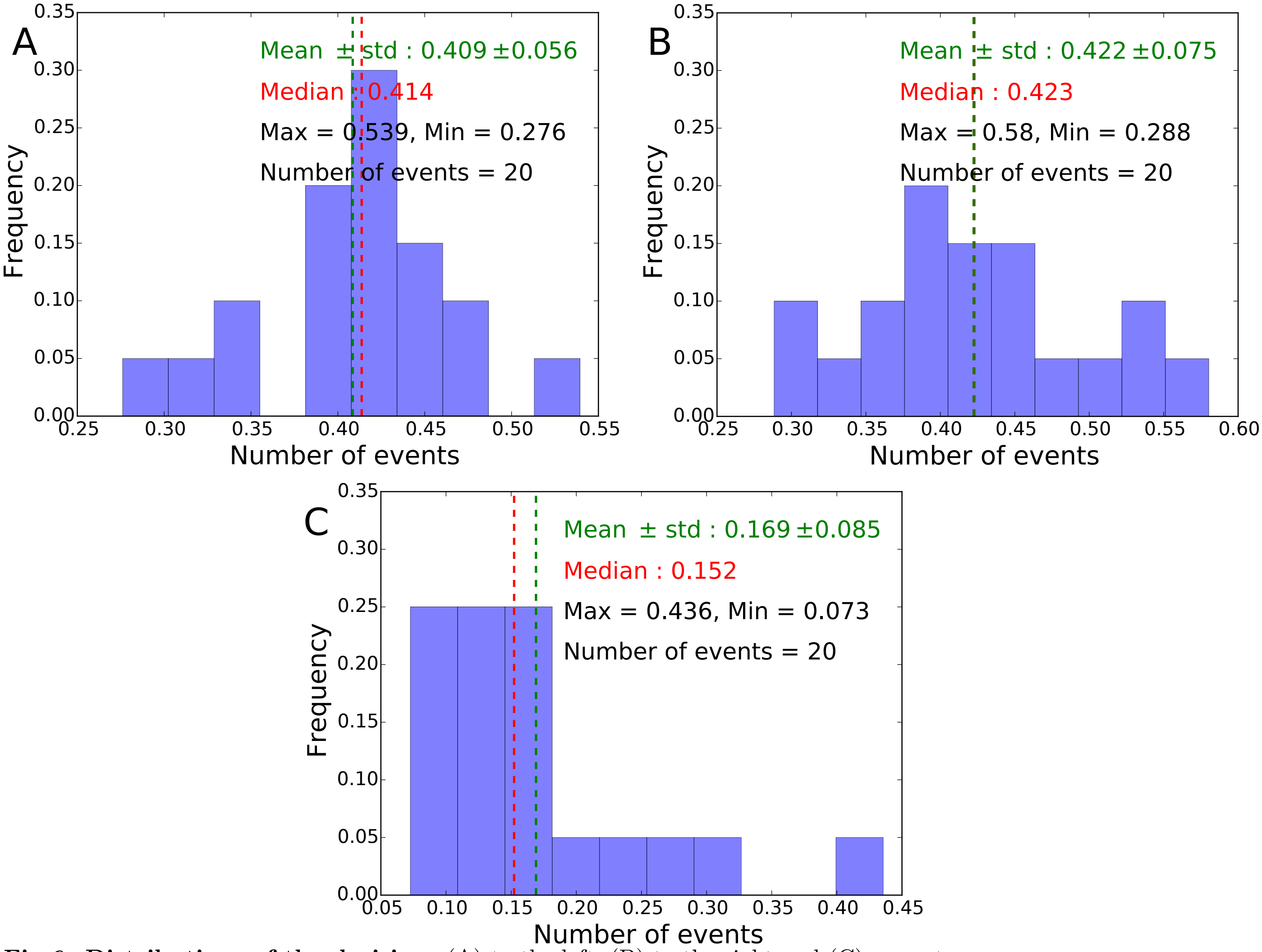


**Fig 6. Distributions of the decisions** (A) to the left, (B) to the right and (C) as a uturn.

#### 1.3 Leadership

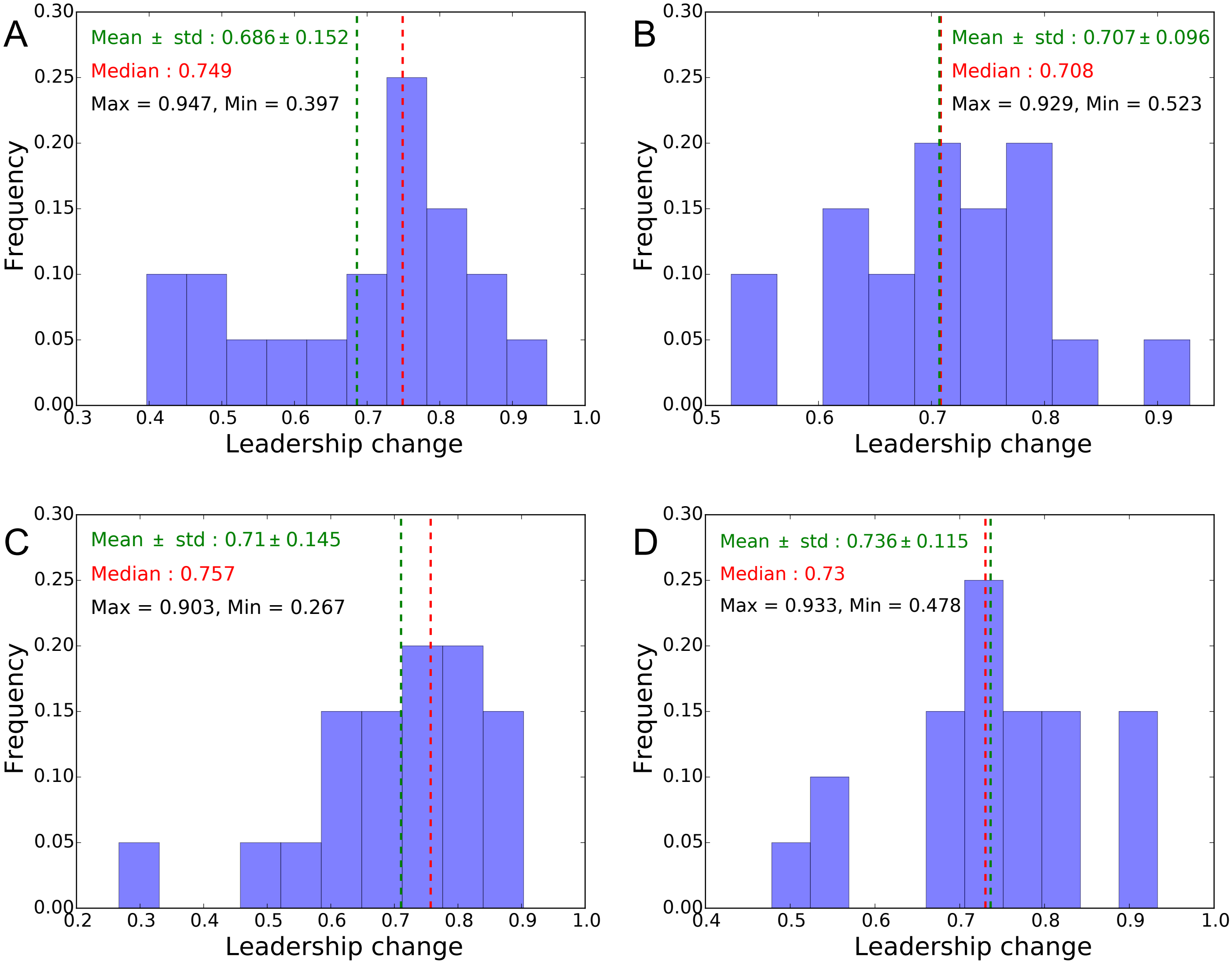


**Fig 7. Distributions of the changes of leadership**. (A) For the interval 0 to 15 minutes, (B) for the interval 15 to 30 minutes, (C) for the interval 30 to 45 minutes, (D) for the interval 45 to 60 minutes.

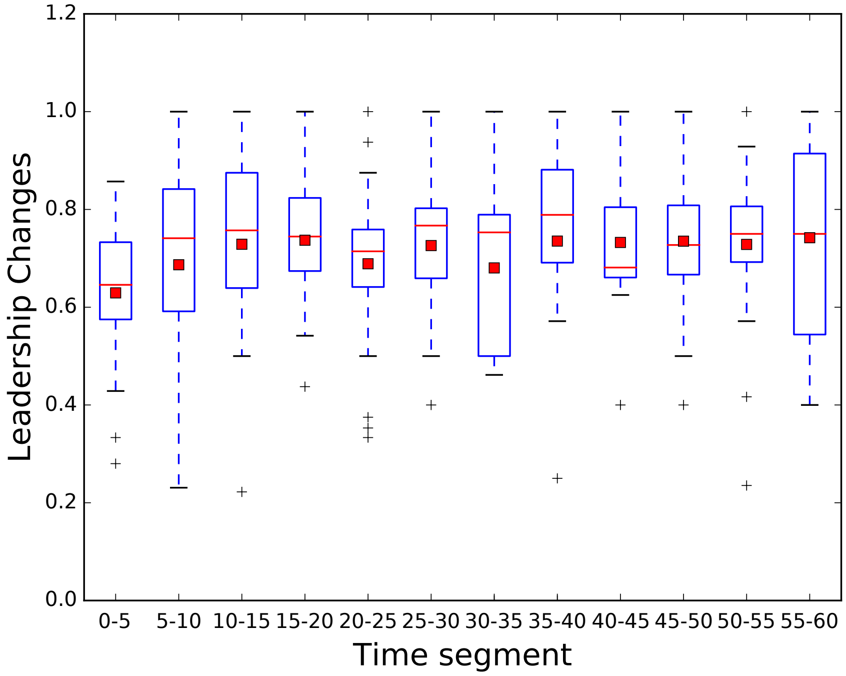


**Fig 8. The leadership changes** at each transition with a high probability and during the whole duration of each replicate. We compared the changes of leadership between consecutive events of decision in the triangle of decision. The red line shows the median and the red square is the mean.

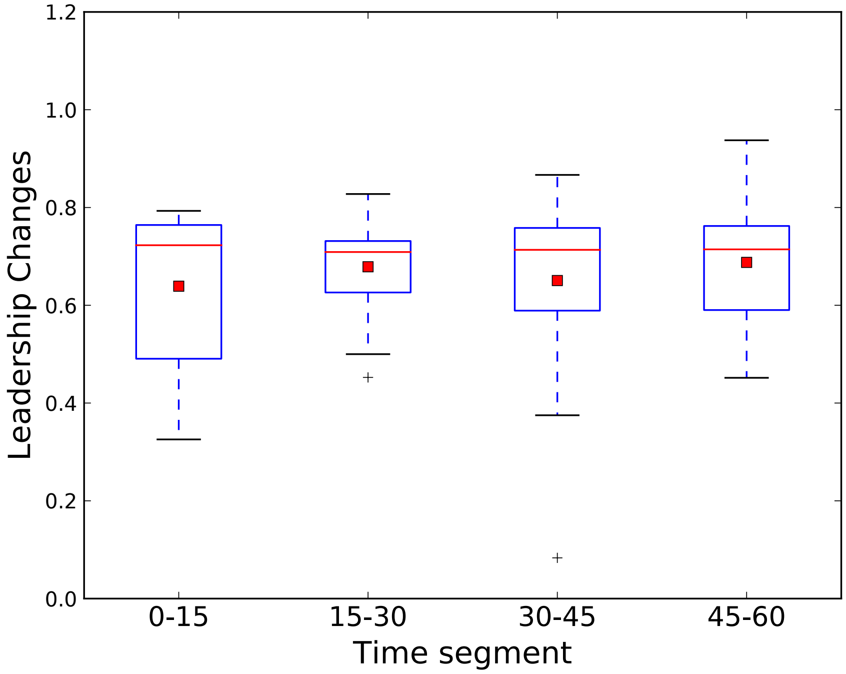


**Fig 9. The leadership changes** at each transition with a high probability and during the whole duration of each replicate. We compared the changes of leadership between consecutive events of decision at the exit of a room. The red line shows the median and the red square is the mean.

The Figure S 10 shows that the number of successful collective departures is proportional to the number of attempts. We considere as a successful collective departure an event when all the population moves from a room to another one. The attempts are all the events where at least a fish leaves a room followed or not by its conspecifics.

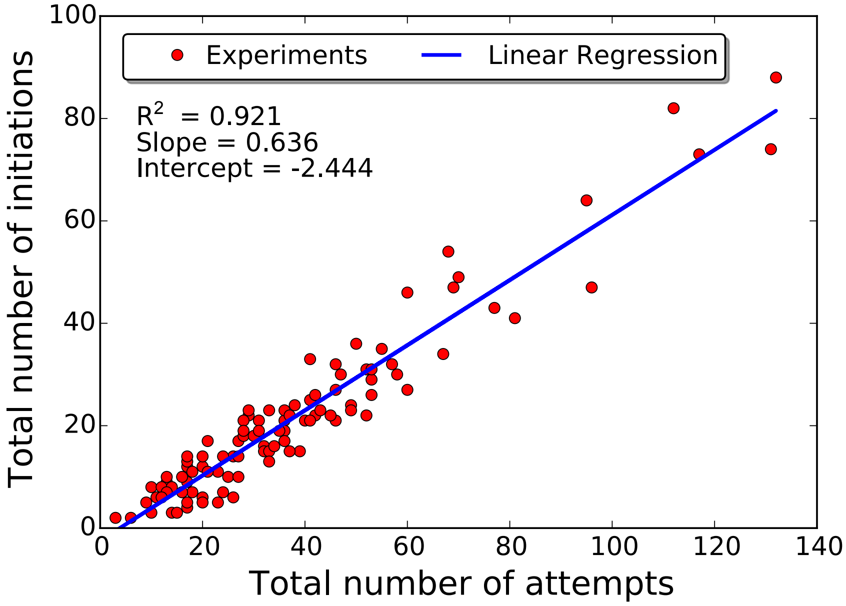


**Fig 10. Successful collective departures and collective departure attempts of the initiator**. The number of successful collective departures is proportional to the number of attempts.

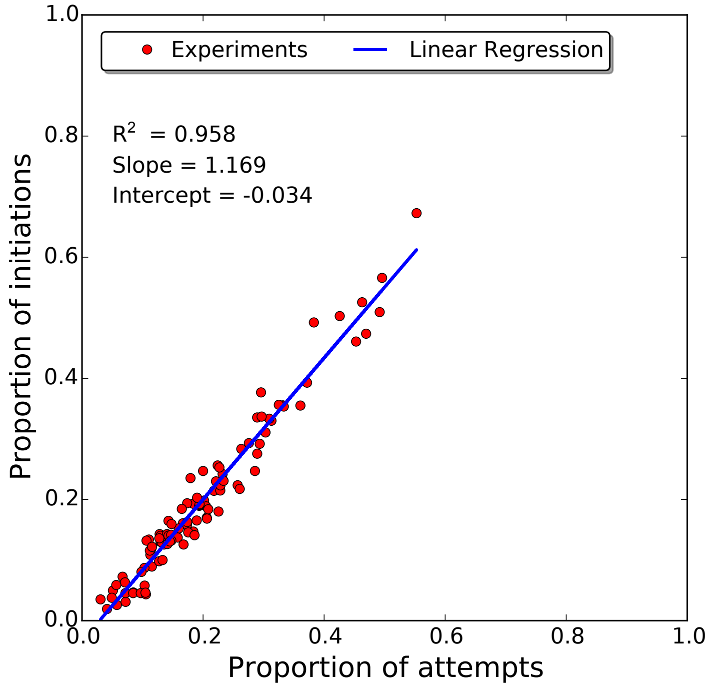


**Fig 11. Successful collective departures and collective departure attempts of the initiator**. Proportion of initiations related to the proportion of attempts.

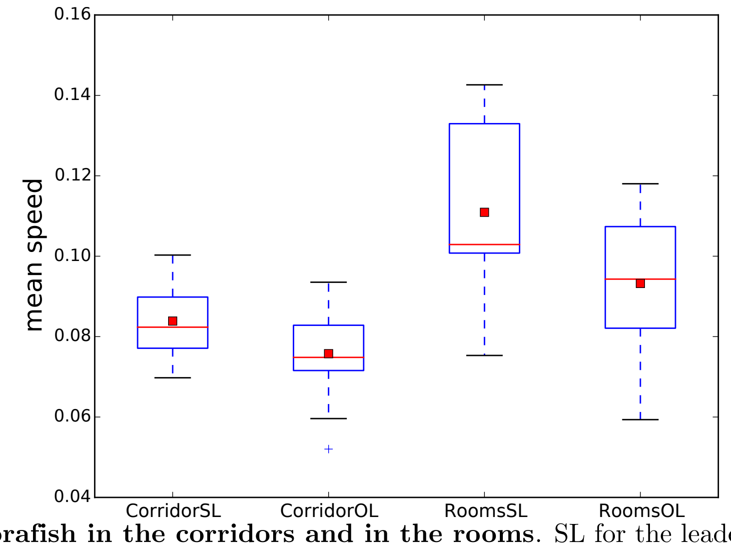


**Fig 12. Speed of the zebrafish in the corridors and in the rooms**. SL for the leaders

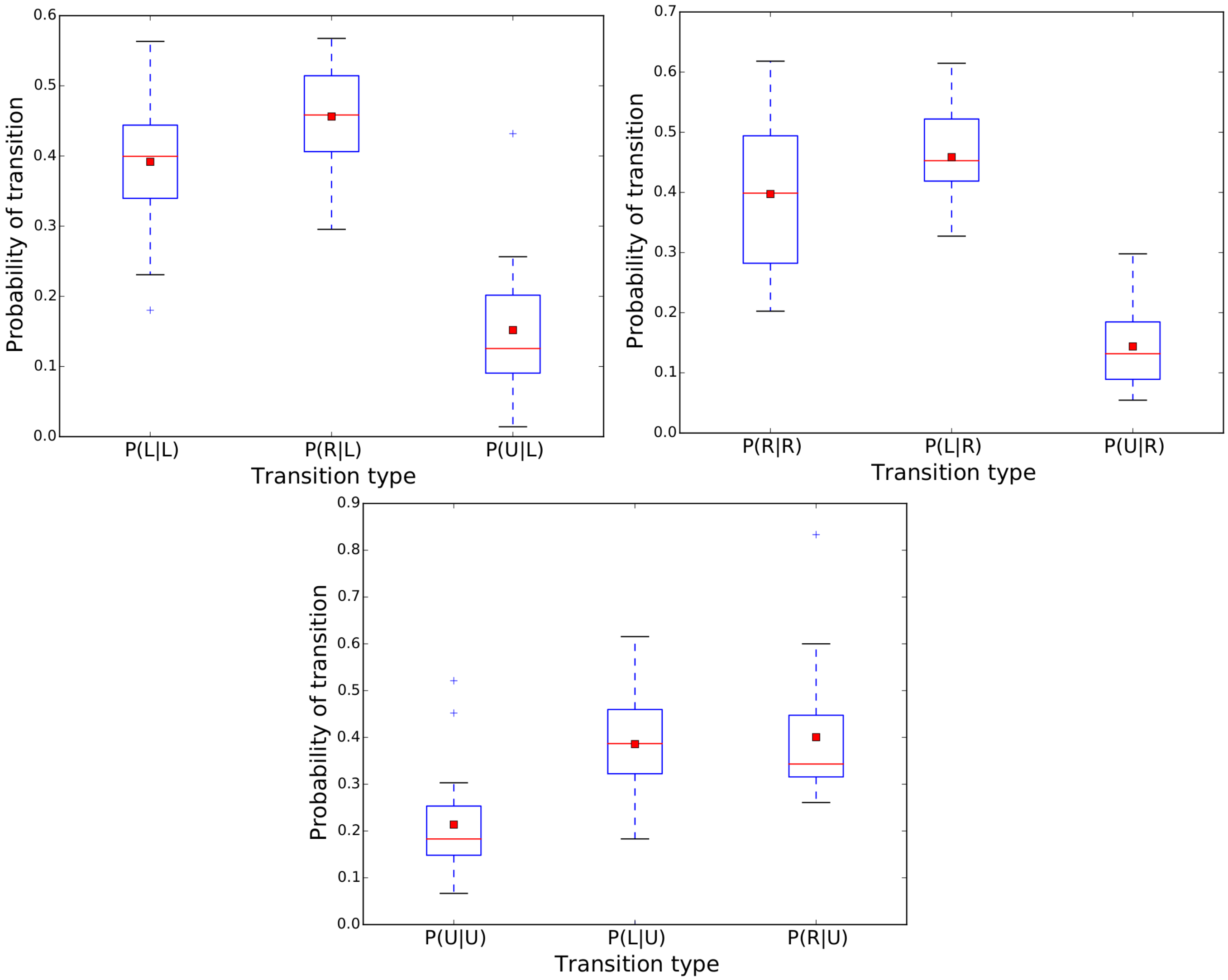


**Fig 13. Probability of the transitions related to the previous transition**. For example, $P(L|L)$ shows the probability $(P)$ for the group to turn to the left $(L)$ when crossing the triangle of decision knowing that $(|)$ the previous transition was to the left $(L)$. $L$ for turning left, $R$ for turning right and $U$ for U-turn.
